# Topographic Connectivity in a Duration Selective Cortico-Cerebellar Network

**DOI:** 10.1101/2020.04.08.031385

**Authors:** Foteini Protopapa, Masamichi J. Hayashi, Ryota Kanai, Domenica Bueti

## Abstract

How does the human brain represent millisecond unit of time? A recent neuroimaging study revealed the existence in the human premotor cortex of a topographic representation of time i.e., neuronal units selectively responsive to specific durations and topographically organized on the cortical surface. By using high resolution functional Magnetic Resonance Images here, we go futher this previous work, showing duration preferences across a wide network of cortical and subcortical brain areas: from cerebellum to primary visual, parietal, premotor and prefrontal cortices. Most importantly, we identify the functional connectivity structure between these different brain areas and their duration selective neural units. The results highlight the role of the cerebellum as the network hub and that of medial premotor cortex as the final stage of duration recognition. Interstingly, when a specific duration is presented, only the communication between the units selective to that duration become particularly “active”. These findings identify duration tuning and topographic connectivity as possible mechanisms underlying our capacity of telling time.

## Introduction

The appreciation of a melody is deeply dependent on the musical tempo i.e., the time with which each single note is played. If this tempo changes, the melody itself changes. Appreciating and playing music requires the very rapid processing of multiple durations. How the brain, encodes and reads out all these different durations remains unclear. A growing body of evidence highlights the contribution of many different brain regions, including cortical (inferior parietal, premotor and prefrontal cortices) and subcortical brain structures to temporal computations. However how these different “time-related” regions communicate with each other and how they contribute to time processing and perception remains largely unknown^1,2^.

In a recent fMRI study^3^, we measured human brain activity at high spatial precision (using a 7 Tesla magnet) while asking participants to discriminate visual stimuli of different durations. In different experimental trials participants were asked to discriminate pairs of durations belonging to different ranges (i.e., from 0.2 to 1 second). The aim of the study was to measure brain responses at the offset of the first stimulus of each pair (i.e., the end of a “purely” encoding stage of the task) to assess the existence of chronomaps i.e., neuronal units selective to different durations and orderly mapped in contiguous portions of the cortical surface. Results revealed the existence in the medial premotor cortex (i.e., Supplementary Motor Area, SMA) and intraparietal sulcus (IPS) of neural units displaying selective responses to stimulus duration (i.e., duration selective clusters of voxels). In these clusters, the hemodynamic response was enhanced by durations in close temporal proximity to the preferred durations and suppressed by durations far from the preferred one. However, only in SMA duration selectivity was organized into topographic maps (i.e., chronomaps). The maps were topographic in that voxels with similar response preference were spatially contiguous.

In humans, topographic maps of stimulus duration and event frequency have been also very recently described not in a single brain region, but in ten different cortical locations, from lateral occipital to inferior parietal and premotor regions^3^. However, in this very recent work, the lack of a duration and/or frequency-related task, make difficult to link these newly discovered maps to duration and temporal frequency perception.

The existence of duration preferences and topography is an intriguing finding, because it suggests that the brain has a seemingly discrete representation of stimulus duration. Additionally, the presence of duration tuning in several cortical areas suggest that this representation is redundant across the brain. To shed light on the tuning mechanism itself and on its redundancy, we decided to use part of the same set of high-resolution fMRI data we used to uncover SMA chronomaps, to answer two additional questions. First, how widespread duration preferences are in the human brain? This question is important since here, differently from what has been recently done^4^, we assessed duration preferences in a duration discrimination task. Second and most important, how do duration-selective units communicate and coordinate with each other? And by answering this last question, our goal is to gain insights on the functional role of the different “time sensitive” brain areas.

In this study we focused on the brain regions that in the original work^3^ were significantly active at the offset of the different encoded durations. These areas were: cerebellum, primary visual cortex (V1), IPS, SMA and Inferior Frontal Gyrus (IFG). In these locations we first assessed the existence of duration preferences and then we checked the connectivity structure between them and between their four “duration channels” (i.e., duration selective clusters of voxels). To this last purpose we measured effective connectivity with Dynamic Causal Modelling (DCM)^4,5^. With DCM we tested various causal models of the interactions between the different network nodes and uncovered the most probable one. Importantly, DCM allowed us to make inferences about the directionality of the connectivity between the different brain areas^4^.

## Results

In the first fMRI experiment of the original work (Exp.1)^6^, we measured brain activity while asking participants (N=11) to decide whether the second stimulus (S2) of a pair, was longer or shorter than the first one (S1, see Fig 1a). S1 could be equal to one of four different duration ranges (0.2, 0.4, 0.6 and 1.0 s) and S2 shorter or longer by a Weber fraction of 0.4. Stimuli were visual gratings (i.e., Gabor patches) varying in both orientation and duration (Figure 1a). Orientation changes were task irrelevant (see Materials and Methods section for details).

**Figure 1.**
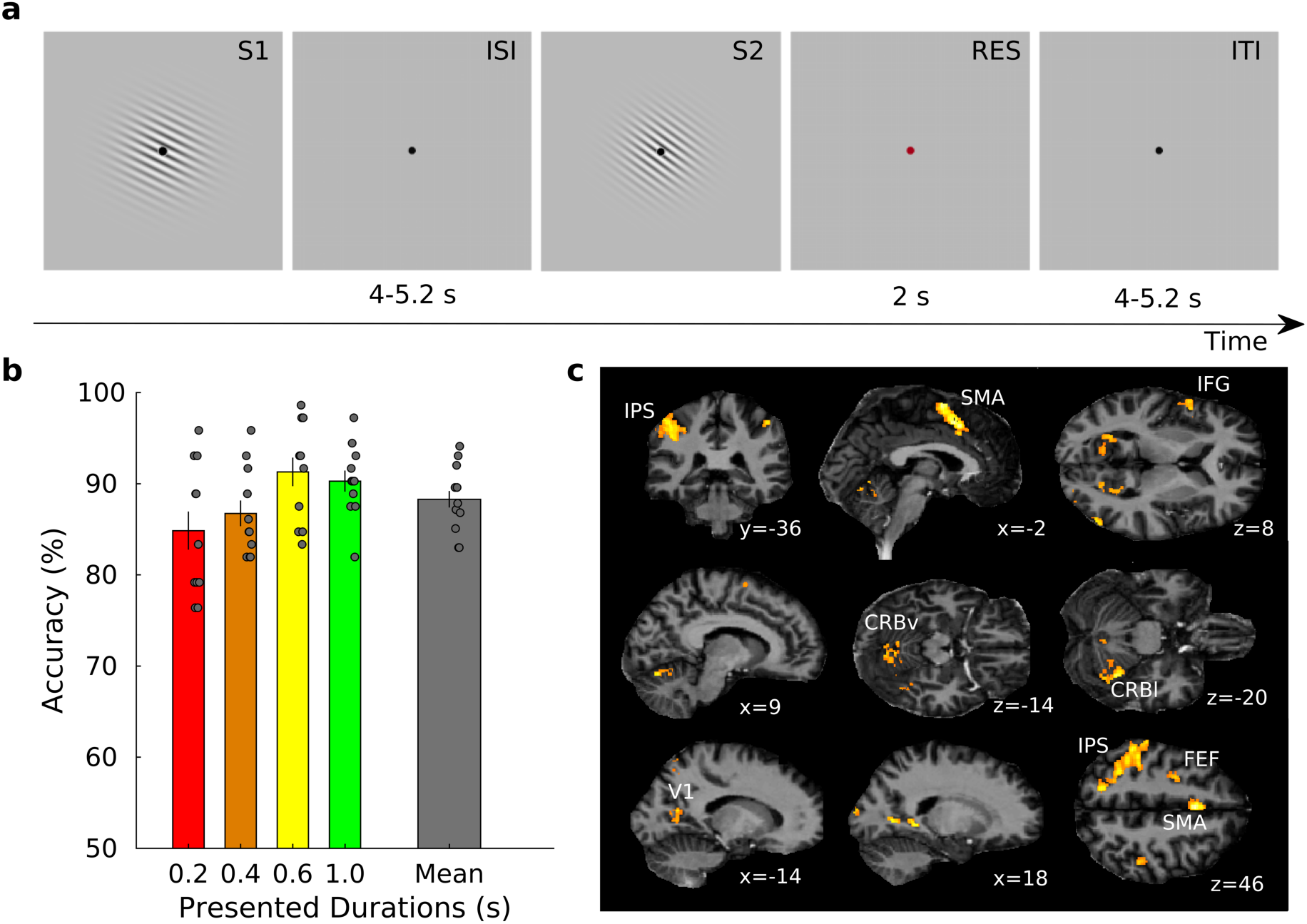
Stimulus sequence, behavioral results and fMRI group results. (a) Schematic representation of the stimulus sequence. In each trial a standard (S1) and a comparison duration (S2) were presented. S1 could be one of four different durations (0.2, 0.4, 0.6, and 1 s) while S2 was either shorter or longer by a Weber fraction of 0.4. Stimuli were sinusoidal Gabor patches varying in orientation. Orientation changes were task irrelevant. Participants were asked, by pressing one of two response keys, to judge whether the duration of S2 was shorter or longer than S1. Inter-stimulus interval (ISI) and inter-trial interval (ITI) varied from 4 to 5.2 s with steps of 0.08 s. Response (RES) period was hold fixed at 2 s. (b) Group average (N=11) of percentage of accuracy plotted separately for each of the four durations and as a mean of them (grey rightmost bar). The grey dots represent the different subjects. Error bars are standard errors. (c) Activations correlated with the offset of the four different S1 durations (p<0.05, FWE cluster level corrected for multiple comparisons across the whole brain). The significant clusters are overlaid on a structural image normalized to the common DARTEL space (see Supplementary Table 1 for more details).

Behavioral data show that participants performed equally well in all four tested durations (see Fig 1b). Proportion of correct responses were 85.1 ± 7.1 (mean ± standard deviation), 87.0 ± 4.9, 91.5 ± 5.4 and 90.6 ± 4.1 % for each S1 duration condition (i.e., 0.2, 0.4, 0.6, and 1.0 s) respectively. Overall accuracy was 88.6 ± 3.7 %. A one-way repeated measures ANOVA with within-subject factor of S1 durations showed a significant main effect (F_3,30_ = 4.824, p < 0.05). Albeit, pair-wise post-hoc tests showed no significant difference between the different combinations of S1 durations (all p’s > 0.05, Bonferroni-corrected for multiple comparisons).

As in the original experiment, also here we focused on the brain response associated with the encoding of the first stimulus of the pair (S1) and specifically the offset of it. We decided to focus on the stimulus offset because this was the moment when the duration of the stimulus became available to the participant. We employed a mass-univariate General Linear Model (GLM) approach at both single subject and group level and used separate regressors for each of the four different S1 durations. These regressors of interest were convolved with the canonical hemodynamic response function (HRF, see Methods section for more details about the modeling of regressors of no interest). We first identified the brain regions significantly associated with the presentation of the four S1 durations together (see Fig 1c and Supplementary Table 1) and then we searched for brain areas that were maximally responsive to each of the four S1 durations separately. For this purpose, we performed a winner-take all classification on the four t-statistics maps, one for each S1 duration, that were generated at both single subjects and group level (see Materials and Methods section for more details on the GLM analysis). The brain areas responsive to the four S1 durations were Cerebellum (Vermis VI, left lobule VI, right lobule V), left and right primary visual cortex (V1), left Intraparietal Sulcus (IPS), left Supplementary Motor Area (SMA) and left Inferior Frontal Gyrus (IFG, see Figure 1c and Supplementary Table 1).

To the purpose of investigating the existence of duration preferences in a wide brain network, we first checked whether each duration selective cluster of the five areas (i.e. from now on called Regions of Interest -ROIs) showed a form of “duration tuning”. From the original work, we knew that both IPS and SMA exhibited duration tuning. However, since cerebellum, V1 and IFG did not show any sign of topographical arrangement of duration selective clusters, duration tuning was not previously assessed in these three regions.

To examine response tuning, for each duration selective cluster of voxels within each ROI, we looked at the changes of the hemodynamic response for the non-preferred durations (2nd brain volume after S1 offset). For each subject, in each duration selective cluster, the hemodynamic response was normalized to the preferred duration, i.e., the duration to which the cluster was maximally responsive to, based on t-statistics maps. As shown in Figure 2b, in all ROIs and all duration selective clusters (i.e., colored lines), the hemodynamic activity was modulated by the S1 durations. Specifically, the hemodynamic response, which as expected peaked at the preferred duration, (PD, see the diamonds in the plot), slowly decayed for durations distant from the preferred one (for all ROIs PD vs PD±1 and PD vs PD±2, p<0.001, see Figure 2c). The gradual decay of the hemodynamic response for durations temporally distant (i.e., non-preferred) from the preferred ones is the most interesting results here, indexing the presence of duration tuning.

**Figure 2.**
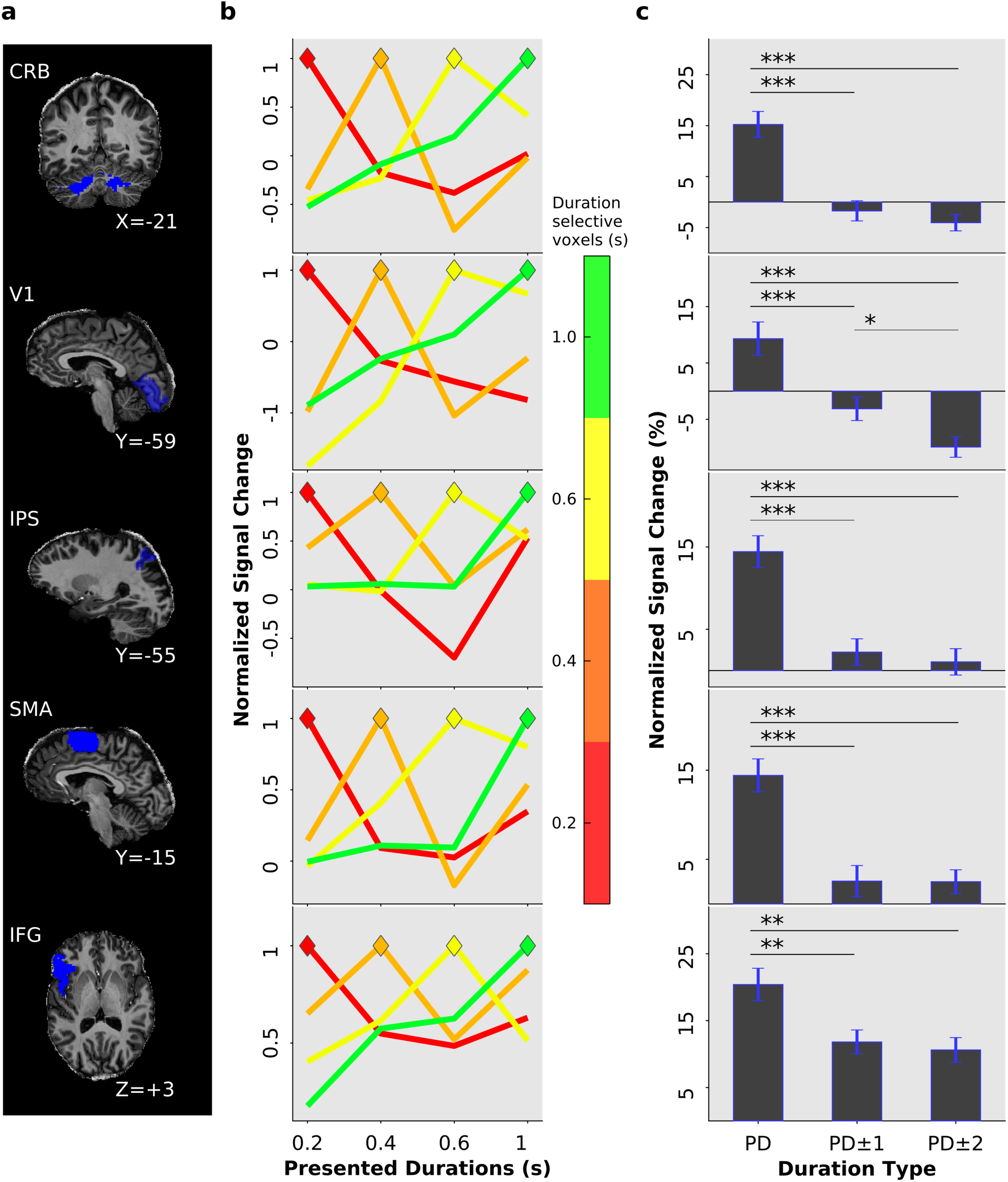
Duration tuning across multiple brain areas. (a) Example of volumetric masks in a single subject. The mask is overlaid on the subject’s native space. (b) Group average of normalized BOLD responses (y-axis) of duration selective voxels (different lines are different clusters of duration selective voxels) for preferred and non-preferred durations. On the x-axis are the four presented durations. The BOLD signal in the duration selective voxels is aligned to the presentation timings of the different S1 durations (i.e., 2^nd^ volume after S1 offset). The colored diamonds represent the point in time where the hemodynamic response of duration selective voxels matched the presentation timing of the appropriate duration (e.g., red-labeled voxels when the shortest S1 duration is presented). The durations of the colorbar are red= 0.2, orange=0.4, yellow=0.6, and green= 1 s and represent the duration-selective voxels. Normalization was performed first in each individual subject to the mean signal intensity across fMRI runs and then for each duration selective cluster to the signal associated to the preferred duration. (c) Normalized BOLD response to preferred (PD), neighboring (PD±1) and distant durations (PD±2) averaged across subjects and duration selective clusters of voxels. Error bars are standard errors. Legend: CRB=cerebellum, V1= primary visual cortex 1, IPS= intraparietal sulcus, SMA=supplementary motor area, IFG= inferior frontal gyrus.

Once we found duration tuning in all five brain areas, we moved to the investigation of the effective connectivity between these areas and their four “duration channels”.

We measured effective connectivity with Dynamic Causal Modelling (DCM). The main objective of DCM is to estimate the coupling parameters of a hypothetical neuronal model and evaluate how well the modeled BOLD signal approximate the observed BOLD response. These coupling parameters concern different aspects of connectivity: the network architecture (i.e., A-matrix) and the influence of a given stimulus (e.g., a visual stimulus lasting 0.2 s) on both the network connectivity (i.e., B-matrix) and the activity of the network node (i.e., C-matrix, see Materials and Methods for details of the DCM equation used). Here, we must stress that the number of possible combinations of these three different aspects of connectivity (i.e., A-B-C matrices) grows exponentially as the number of brain regions included in a system grows. For this reason, rather than exploring the effective connectivity in a 20-nodes network (i.e., the five ROIs by the four “duration channels”), a network for which the number of possible models to test would be exceedingly high, we decided to start with the investigation of the effective connectivity in a simpler 5-nodes network (i.e., the five ROIs without considering the “duration channels”). We also decided to break the whole analysis process in two main stages.

The first set of analyses (for a summary of all performed DCMs see Supplementary Table 2) concerned this 5-nodes network. Our aim here was to explore within this simpler network two features of network connectivity: the network architecture (i.e., how the different network nodes communicate and coordinate with each other, A-matrix) and the influence of a certain stimulus duration on the activity of any of the network node (C-matrix). Once identified these two features, we moved to the second analysis stage and we used these features to inform and therefore simplify the DCM on the 20-nodes network.

Since we did not have a clear hypothesis about the network architecture of the five duration selective brain regions, to identify the connectivity structure of this network we employed a “data-driven” approach, called Parametric Empirical Bayesian method (PEB). Briefly, PEB seeks the optimal model by testing various combinations of A-B-C-parameters through a pruning process. PEB starts with a fully connected and a fully modulated model and then proceeds by switching off certain model’s parameters (e.g. connections) and measures how this affects the model evidence. For PEB analysis we first specified a fully connected model between the five ROIs and we ran the DCM routine of 198 models (a model for each fMRI run and each subject i.e., 11 subjects by 18 runs, see Material and Methods for more details). Models were treated as random effects that could differ between subjects. Posterior probability (Pp) for second-level effect was set to Pp greater than 0.99. According to PEB, the optimal network architecture was the one in which the cerebellum is the only brain area connected with all other network nodes (See Figure 3a). Cerebellum has indeed both feedforward and feedback connections to and from V1 and IFG, and only feedforward connections to SMA and IPS. V1 and IPS have feedforward connections to SMA. IFG has feedback connections from both cerebellum and V1. SMA is the only node of the network that while being affected by the activity of other brain regions, its activity does not affect any other network node.

**Figure 3.**
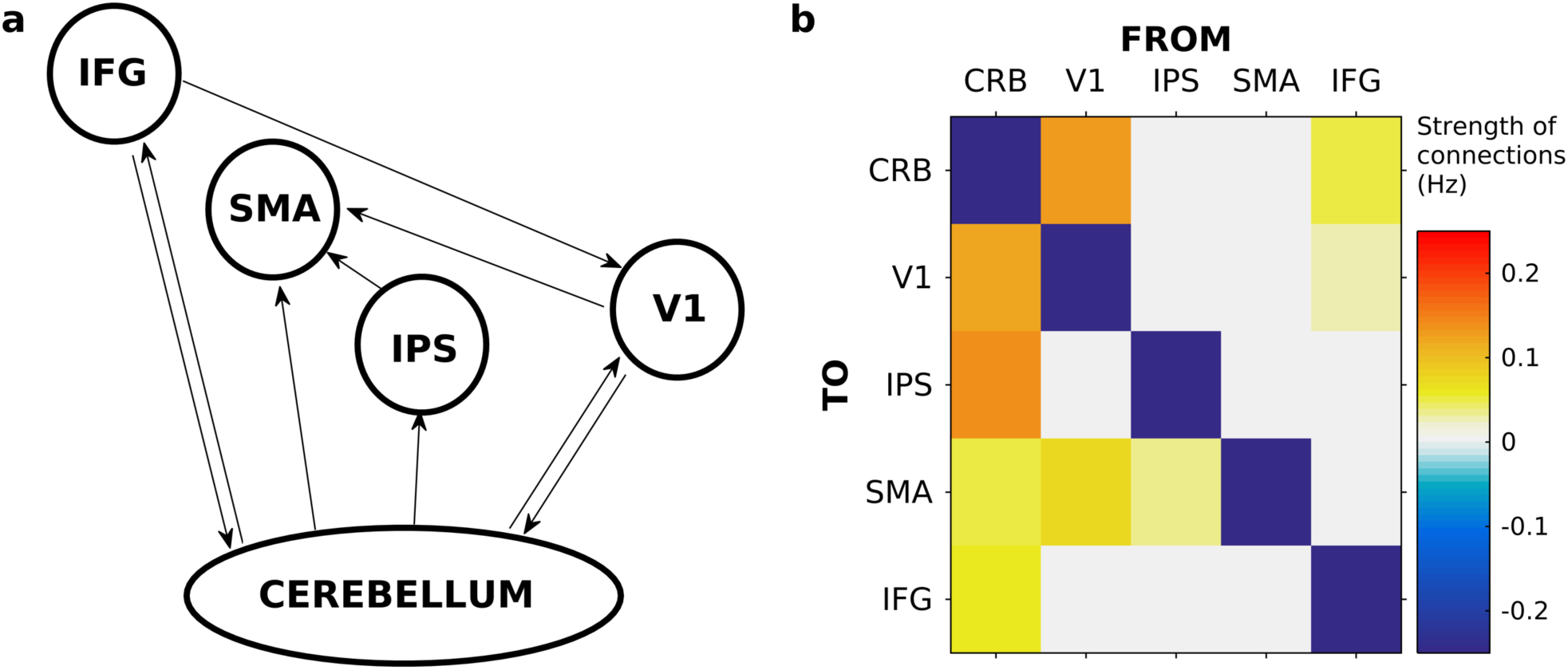
Five-nodes network architecture resulting from Parametric Empirical Bayesian (PEB) analysis. (a) Caricature of the 5-nodes network where the nodes are represented with ellipses and the direction of connections with arrows. (b) Colormap which depicts the strength of the survived connections after the pruning of the parameters done by PEB. Values are in Hz and the range goes from -0.2 to 0.2 (see colorbar). Legend: CRB= cerebellum, V1=primary visual cortex, IPS= intraparietal sulcus, SMA=supplementary motor area, IFG= inferior frontal gyrus.

After the identification of the optimal model across subjects (Figure 3a), we proceeded with Bayesian Model Averaging (BMA, i.e., averaging the parameters across the “winning” models of all subjects), to determine the parameters strength of this connectivity structure (Figure 3b). The values in the matrix depicted in Figure 3b, reflect the strength with which the activation changes in one node affect the interconnected nodes.

To check the robustness of the PEB connectivity results, we ran two additional DCM analyses with two distinct connectivity structures (A-matrix): a “fully connected” and a “PEB-like” version of it. In both DCM models, B and C matrices were left fully modulated. We then compared the two models by using Bayesian Model Selection (BMS)^7^ and we computed their protected exceedance probabilities (PEPs). We considered mandatory for a “winning” model to have PEPs greater than 90%. As shown in Figure 4a the winning model is the one, whose architecture is “PEB-like”. This last DCM analysis validates the results of the initial PEB analysis.

**Figure 4.**
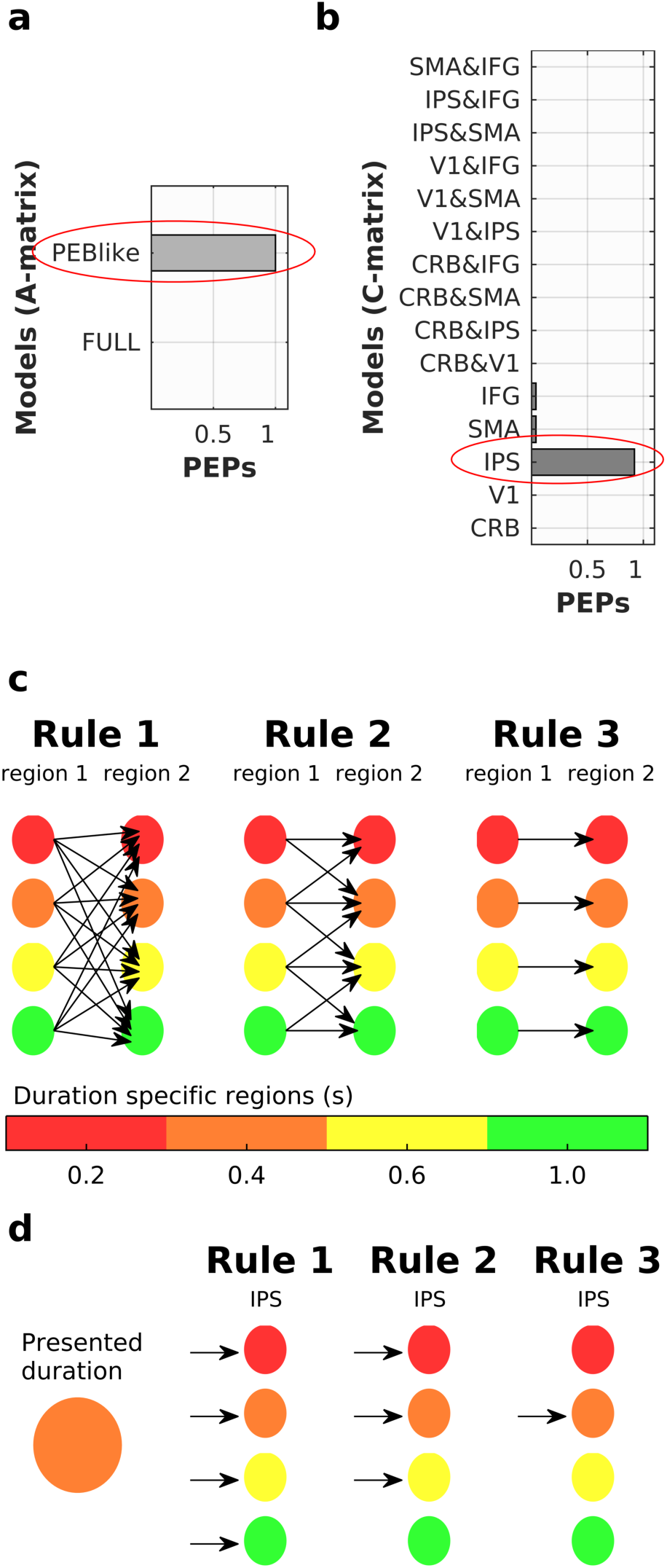
Bayesian Model Selection (BMS) results and rules of connectivity in a 20-nodes network. (a) Bar plot of protected exceedance probabilities (PEPs) of two models with different network architecture (A-matrices) as resulting from BMS. One is the fully connected (FULL) and the other one has a “PEB-like” structure. The red ellipse highlights the winning model. (b) Bar plot of protected exceedance probabilities (PEPs) of the fifteen models that have a “PEB-like” architecture but differ in terms of which region is modulated by duration presentation. The red ellipse highlights the winning model. (c) Rules of connectivity among two hypothetical brain areas. The different duration selective clusters are depicted as color disks. Different colors represent different durations. Rule 1 *duration independent* rule: allows all duration-specific clusters within region 1 to connect to all the clusters within region 2; connectivity with distant sub-regions is allowed (Preferred Duration -PD±2). Rule 2 *neighboring dependent* rule: allows each duration specific cluster within region 1 to connect to the same duration specific cluster in region 2 and its immediate neighbors (PD±1). Rule 3 *duration dependent* rule: allows only the existence of connections among the same duration specific clusters of the two regions (PD). The same rules apply for the modulatory effect that stimulus presentation has on network connectivity and on the activity of the duration selective clusters within the IPS (d). The durations are represented in the colors of the colorbar red= 0.2, orange=0.4, yellow=0.6, and green= 1 s.

After knowing the connectivity structure of a 5-nodes network, we proceed by finding out which region(s) among the five ROIs, was modulated by stimulus presentation (C-matrix). Fifteen possible models with 15 different combinations of C-matrices were tested. In the C-matrices of these 15 models, we arbitrarily allowed to have one or maximum two brain regions modulated by event duration. All models had an A-matrix “PEB-like” and a B-matrix in which all existing connections were modulated by stimulus presentation. Thus 15 possible model variations (i.e., nested models) were constructed using Bayesian Model Reduction (BMR). Bayesian Model Selection (BMS) showed that the most likely model was the one with a C-matrix in which IPS was the area modulated by duration presentation (Figure 4b).

Once identified the best network architecture (A-matrix) between the five ROIs and established that the IPS is the brain area mainly affected by stimulus presentation (C-matrix), we used this knowledge to inform the DCM analysis on the 20-nodes network. In this last analysis we tested specific hypotheses of connectivity between the five ROIs and their four duration selective clusters. These hypotheses concerned the existence of connections (A-matrix “PEB-like”) and the modulation exerted by stimulus duration on both connections (B-matrix) and IPS activity (C-matrix). Specifically, we hypothesize these connections and modulations to be: 1) stimulus *duration independent*, 2) only partially stimulus duration dependent i.e., *neighboring dependent* or 3) totally stimulus *duration dependent* (see Figure 4cd).

We tested 108 possible models which were the combination of these three possibilities (i.e., duration *independent, neighboring dependent, duration dependent*) applied to the A-B and C matrices (see Materials and Methods section for more details). In other words, we tested models with an A-matrix “PEB-like” and where the C-matrix concerned the impact of S1 stimulus duration on the activity of the four duration selective clusters within the IPS.

Although our main interest was to check connections (A-matrix) and modulations (B-matrix) between the different duration-selective clusters across the five ROIs, for completeness we also assessed these connections and modulations within a given ROI.

For the calculation of the A-B-C-parameters of these 108 models we used the Bayesian Model Reduction (BMR) procedure. To identify the winning model, we first divided the 108 models into three main families (36 models each) based on the modulatory effect of S1 stimulus duration on the activity of the IPS duration selective clusters (i.e., stimulus *duration independent, neighboring dependent*, stimulus *duration dependent*, see Materials and Methods for more details) The PEPs of each family of models was then estimated using BMS where models were treated as random effects^8,9^. The winning family was the one in which the presentation of a stimulus duration modulates the IPS activity of the duration selective clusters in a *duration dependent* manner (Figure 5a and c). This means, that the presentation of a 0.4 s stimulus, for example, modulates the activity of the IPS cluster that is selective for that duration only.

**Figure 5.**
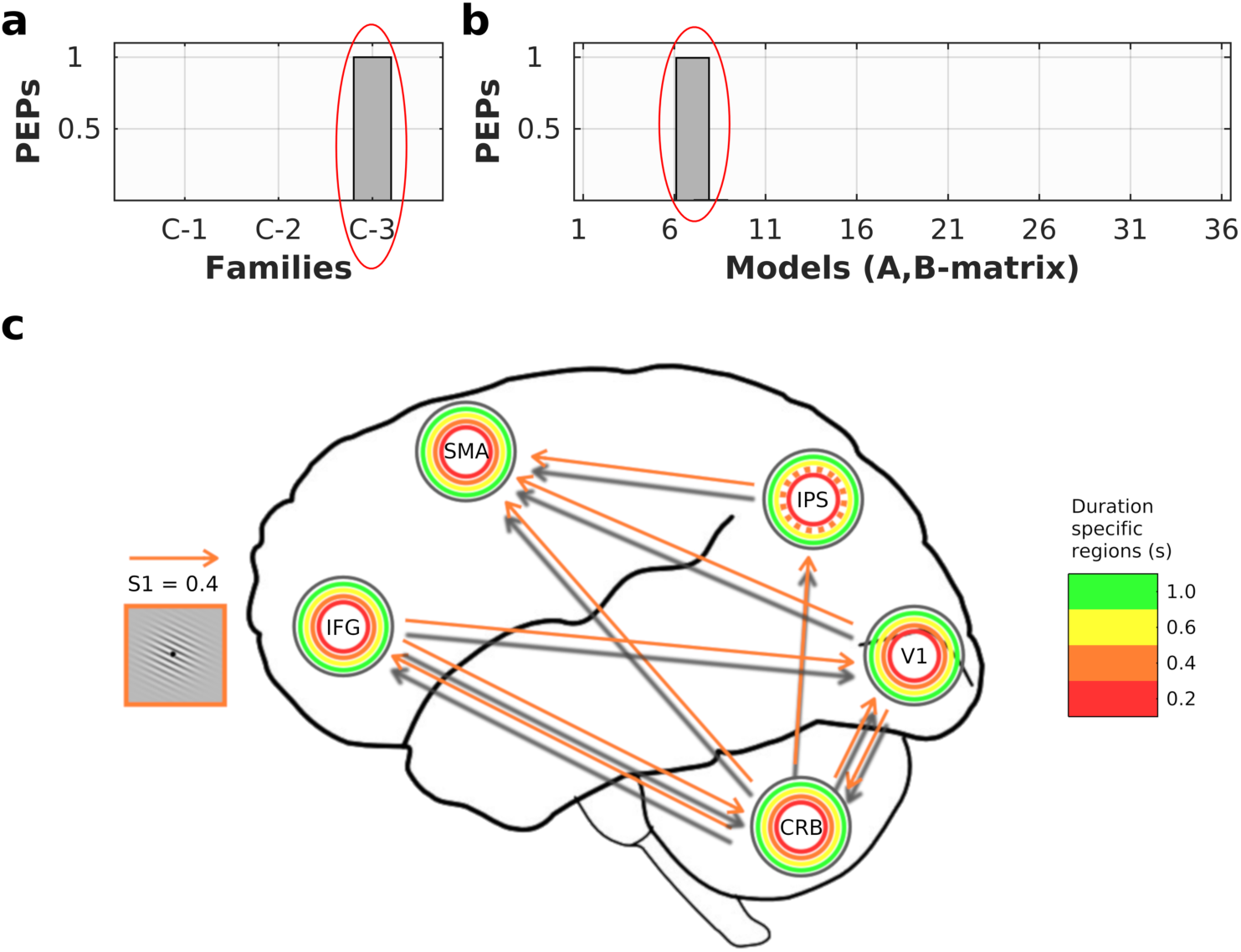
Bayesian model selection (BMS) and Bayesian model comparison results of the 20-nodes network. (a) BMS among three families of models that differ according to the modulatory effect of stimulus duration on the activity of IPS duration selective clusters (i.e., C-matrix that could be: stimulus *duration independent, neighboring dependent*, or stimulus *duration dependent*). Bar plot shows their protected exceedance probabilities (PEPs). Each family consists of 36 models. The red ellipse highlights the winning family i.e., family of models where the modulation by S1 stimulus duration on IPS activity is stimulus *duration dependent*. (b) Bayesian model comparison (BMC) across the 36 models of the winning family. Bar plot depicts the protected exceedance probabilities (PEPs). The 36 models have C-matrix stimulus *duration dependent* and an A and B matrix “PEB-like” that could be stimulus *duration independent, neighboring dependent*, or stimulus *duration dependent* within and between ROIs. The red ellipse highlights the winning model. (c) Schematic representation of the winning model on a brain silhouette. The network architecture “PEB-like” (A-matrix) is represented in the arrows’ shadow (gray arrows). The five different ROIs are the different grey disks, whereas the duration selective clusters are the embedded colored disks with different diameter. The *duration dependent* modulation of S1 stimulus presentation on IPS activity (C-matrix) is represented in the dotted border of the orange duration selective cluster. The *duration dependent* modulation on the “PEB-like” network connectivity (B-matrix) is represented by the orange arrows.

We then determined which of the 36 models within the winning family of models (where activity of IPS was modulated in a duration dependent fashion), best explained our data. We applied Bayesian Model Comparison (BMC)^9^, to compare models differing in connectivity (A-matrix) and modulation of connectivity by stimulus duration (B-matrix) according to the different levels of duration specificity (see Figure 4c). A caricature of the winning model is depicted in Figure 5c. In this model all hypothesized connections resulting from the PEB analysis are present (A-matrix “PEB-like” fully connected, see Figures 3a and 5c grey arrows). However, when a specific duration is presented, e.g. 0.4 s, there is a change in the connectivity that is *duration dependent*, i.e., it affects the connectivity between the 0.4 duration selective clusters of the functionally distinct brain areas (e.g., the 0.4 cluster in cerebellum and V1 see the orange arrows in Figure 5c). Within each brain region though, this modulation is duration *independent*, for example the activity of all duration selective clusters within the cerebellum are influenced by the presentation of a 0.4 s duration. Finally, to check the parameters of the winning model across subjects, we use Bayesian Parameter Averaging (BPA) to average the A-B-C-parameters of the winning model across subjects^10^ (See supplementary Figure 1).

## Discussion

In this study we used fMRI at ultra-high field to show the existence in a wide circuit of brain areas, including cerebellum, primary visual cortex, intraparietal sulcus, supplementary motor area and inferior frontal gyrus, of neuronal units (clusters of voxels) displaying duration preferences. In each duration selective cluster of voxels the signal amplitude decreased for durations distant from the preferred one. When we looked at the connectivity architecture of these five functionally distinct brain areas (PEB analysis on a 5-nodes network), we found that the optimal model is the one in which cerebellum has feedback and/or feedforward connections from and to all other network nodes. Whereas SMA is the only area that, while being modulated by the activity of cerebellum, IPS and V1, does not influence the activity of any other brain region. Moreover, we found that within this network, the brain area greatly affected by stimulus presentation was the IPS. We then explored the effective connectivity between these five regions and their four duration selective clusters of voxels. The DCM analysis of this 20-nodes network, showed that while all connections between the nodes exist in the form resulting from the PEB analysis, the connectivity of the duration selective clusters across functionally distinct brain regions was modulated in a *duration dependent* fashion. In the same vein, the modulation of stimulus presentation on the activity of the four IPS duration selective clusters was also duration specific.

The existence of duration tuning across different brain regions is an intriguing and novel finding. Electrophysiological studies in animals, have shown the existence in numerous brain areas of cells selectively responsive to duration and intervals on the order of tens-to-hundreds of milliseconds^11,12,13^.However, only very recently duration preferences have been described in the human brain^6,3,14^. In a recent fMRI study, for example, Harvey and colleagues showed the existence of a topographic representation of event duration and frequency in ten cortical locations along a functional hierarchy that goes from occipital to frontal regions^3^. Maps were defined as changes in the response preference of voxels to duration and frequency as a function of cortical distance i.e., voxels with similar response preference were spatially contiguous. However, in this study, for most of the trials, subjects passively looked at the stimuli and only occasionally were asked to detect a color change in them. It is therefore unclear how activity in these duration and frequency maps relates to duration and frequency perception. Here instead, we report duration tuning in a network of brain areas that were significantly associated with duration encoding in a duration discrimination task. It is thus likely that both the tuning and the network connectivity shown here are linked to aspects of duration perception and discrimination. The existence in the brain of discrete, separable representations of time is fascinating. This seemingly discrete representation, likewise to the representation of other sensory features, for example shape, speed, pitch, serves very likely the purpose of improving the identification and the discriminability of different durations^15,16,17^. This may be very important in situations where a rapid and accurate processing of a multitude of different durations is needed, for instance, in playing music or understanding language. Why these specific representations of duration are redundant across brain areas is unclear. A possibility is that despite this similar representation of time, cerebellum, V1, IPS, SMA and IFG play a different role in duration encoding and perception.

In the direction of finding out the role played by these different brain regions in duration perception, the connectivity results are of a great help. PEB results indeed seem to suggest the existence of specific functional relationships between the network of areas active at duration encoding. Specifically, the cerebellum seems to work as network hub. With its feed-forward and feedback connections to and from all the other network nodes, the cerebellum seems in charge of both controlling the flow of information and updating the state of the network. Interestingly, the only bidirectional connections that the cerebellum has, are with V1 and IFG, i.e., the two edges of a hypothetical functional hierarchy. V1 is probably the sensory input area, the area conveying the temporal signal embedded in the visual input. Whereas the IFG could be the area where encoding-related decisions takes place, as previous studies have shown^18^. Both V1 and IFG receive and send information to cerebellum, but the direct connection between IFG and V1 is unidirectional only i.e., IFG modulates V1 activity and not vice versa.

Cerebellum has also feedforward connections to IPS and SMA and these connections are unidirectional. This last result seems to suggest once again a key role of the cerebellum in mediating the temporal encoding process. A possibility is that the cerebellum mediates the flow of information between input and output regions and in doing so talks to core timing structure such as IPS and SMA. Since IPS is the only brain area whose activity is greatly affected by duration offset, it can be considered the place where a first reading of temporal signals occurs (i.e., “duration input” area). SMA instead, being modulated by the activity of both IPS and V1, but not modulating the activity of any other network node, could be considered the final stage of duration recognition. In humans, the role of both SMA and inferior parietal lobule in temporal perception has been extensively documented^19^. Both areas have been implicated in a variety of timing tasks^20,21^ with a range of durations spanning from a few hundreds of milliseconds to a few seconds^22,23^ and with stimuli of different sensory modalities^24,25^. It is therefore likely that both areas constitute a core of the timing network. The functional link between SMA and inferior parietal lobule activity has been recently shown by mean of Transcranial Magnetic Stimulation (TMS) combined with electroencephalography^26^. In this study, the perturbation of the right supramarginal gyrus (SMG) just before the encoding of a stimulus duration, affects activity in frontocentral sites including SMA. These effects correlate with changes in duration perception, suggesting thus a direct link of SMG and SMA connections with duration perception. Also cerebellum has been proved to play a key role in motor and perceptual timing tasks of discrete isolated intervals^27,28,29^. Specifically, the cerebellar lobule V and VI, where we observed activity correlated to duration encoding, have been previously associated to interval timing^30,31^, motor and working memory tasks^32^. These lobules, according to resting-state connectivity maps, are part of somatomotor, ventral attention and frontoparietal networks^33^. The computational mechanism through which cerebellum contributes to millisecond time perception is still a matter of debate. Our data here seem to suggest a role of hub of the cerebellum, coordinating the input (V1) and output (IFG) regions by talking with core timing areas (SMA and IPS).

When we considered the five network nodes and their four durations selective clusters, we found that at stimulus offset, the full connectivity pattern described by the PEB analysis was present. However, this pattern was modulated across regions in a *duration dependent* manner. In a similar vein, also IPS activity was affected by S1 duration offset in a duration specific fashion. Both these results are compatible with the presence of tuning in all these five brain areas. When a given duration is presented, it makes sense that across the five nodes, only the communication across appropriate “duration channels” become particularly “active”. This result likely serves the efficiency of the neural communication^17^. A one to one communication indeed, enables information entering at the input level to be reproduced faithfully at the output level. And having a more efficient communication saves metabolic resources and time^17,34^. Our hypothesis is that this topographic connectivity i.e., a one to one communication between duration selective clusters, either supports or is a consequence of the tuning mechanism.

Due to the limited spatial resolution of fMRI, it is impossible to draw a direct link between the duration preferences of the clusters of voxels in humans, with the duration tuned cells in animals^12,35^. It is indeed difficult to establish whether the changes detected at voxel level reflect the activity of different populations of neurons tuned to different durations or whether it is the same population whose neuronal dynamics change depending on the presented duration^36,37^. What we can conclude based on our findings is that time perception and encoding is the result of the activity of a network of brain areas and it seems supported by duration tuning mechanisms and topography at both representation and connectivity level.

## Materials and Methods

The data of this study are partially shared with two published studies^6,38^

### Participants

We tested eleven healthy volunteers (5 females, mean age 23.7 years, SD 4.3 years) with normal or corrected-to-normal vision. All volunteers gave written informed consent to participate in the study. The experimental protocol was approved by the ethics committee of the Faculty of Biology and Medicine at the University Hospital of Lausanne (protocol number 92/2012) in accordance with the Declaration of Helsinki.

### Stimuli and Procedure

The volunteers performed a temporal discrimination task of visual durations. Visual stimuli were sinusoidal Gabor patches (100% contrast, spatial frequency of 1.9 cycles/degree, Gaussian envelope with standard deviation of 2.2 degrees, diameter of ∼9 degree) with a central circular hole (diameter 0.6 degrees) displayed at the center of the screen around a central fixation point (a black disk 0.5 degrees of diameter at a viewing distance of 90 cm) on a grey background.

In each trial, two Gabor patches (S1 and S2) were sequentially presented at a variable inter-stimulus-interval ranging between 4 and 5.2 s (in 0.08 s steps). The two stimuli were followed by a response cue (a red fixation point) of 2 s duration (see Figure 1a). S1 and S2 varied in both orientation and duration but only duration was task relevant. The duration of S1 could be 0.2, 0.4, 0.6, and 1 s and its orientation 36, 72, 108, and 144 degrees. S2 could be either shorter or longer than S1 by a constant Weber fraction of 0.4. The orientation of S2 was a value randomly chosen from the four possible orientations used for S1. Participants were asked to decide whether the duration of S2 was shorter or longer than S1 by pressing one of two buttons on a response-pad. The combination of different durations and orientations lead to 16 different trial types. Each S1 stimulus type was presented only once in each fMRI run. A total of 18 fMRI runs were collected in two separate sessions (9 runs per session with 1-3 days of latency). The total duration of each run was 3 min and 51 s.

### Behavioral Data Analysis

A one-way repeated measures ANOVA was carried out on percentage accuracy values of each individual subject. The alpha level was set to 0.05, while Bonferroni test was used as post-hoc test.

### MRI Acquisition

Blood oxygenation level-dependent (BOLD) functional imaging was performed using an actively shielded, head-only 7T MRI scanner (Siemens, Germany), equipped with a head gradient-insert (AC84, 80 mT/m max gradient strength; 350 mT/m/s slew rate) and 32-channel receive coil with a tight transmit sleeve (Nova Medical, Massachusetts, USA). The ultra-high magnetic field system allowed us to obtain smaller size voxels thus to increase the spatial resolution of the functional data. This is due to the fact that 7T systems acquire improved BOLD signals; The signal strength of venous blood is reduced due to a shortened relaxation time, restricting activation signals to cortical grey matter. Time-course series of 169 volumes were acquired for each run using the 3D-EPI-CAIPI sequence^39^. The spatial resolution was 2.0 mm isotropic, the volume acquisition time was 1368 ms, the flip angle was 14 degrees, the repetition time (TR) 57 ms and the echo time (TE) 26 ms and the bandwidth 2774 Hz/Px. The matrix size was 106 x 88 x 72, resulting in a field of view of 210 (AP) x 175 (RL) x 144 (FH) mm. An undersampling factor 3 and CAIPIRINHA shift 1 were used. Slices were oriented transversally with the phase-encoding direction left-right. 42×45 reference lines were acquired for the GRAPPA reconstruction. For each subject, a total of 3,042 volumes (169 volumes per run, 18 runs) were analyzed.

High-resolution whole-brain MR images were also obtained using the MP2RAGE pulse sequence optimized for 7T^40^ (voxel size = 1.0 x 1.0 x 1.0 mm, matrix size 256 x 256 x 176, TI_1_/TI_2_=750/2350ms, α_1_/α_2_ = 4/5 degrees, TR_MP2RAGE_/TR/TE = 5500/6.5/2.84 ms).

### fMRI Preprocessing

Functional imaging data were preprocessed using Statistical Parametric Mapping (SPM12 v. 7219, Wellcome Department of Imaging Neuroscience, University College London) toolbox in MATLAB. The EPI volumes acquired in each session were realigned to the mean of the session and then co-registered to the T1-weighted image acquired in the same session. To perform group level analysis the realigned and co-registered images were then normalized to the averaged DARTEL template (Diffeomorphic Anatomical Registration Through Exponentiated Lie algebra)^41^ and smoothed with a 2 mm full-width at half-maximum Gaussian kernel. To perform connectivity analysis, data were also kept in the subject’s native space i.e., data, after realignment and co-registration to the T1-weighted image, were directly smoothed with a 2 mm full-width at half-maximum Gaussian kernel.

### Identification of Regions of Interest (ROIs)

For ROIs identification, we performed a mass-univariate General Linear Model (GLM) approach. The first level analysis was performed on both DARTEL normalized and subject’s native space images. A part from the different space of the images, these analyses were identical.

Each single-subject model included 18 runs/sessions with 6 event-types (model regressors) in each session. Given that we were interested in the encoding of the 4 different durations we used 4 regressors of interest time-locked to the offset of S1 (standard duration), i.e., the moment when the duration of a stimulus became available to participants. We also used 2 more regressors of no-interest; one time-locked to the onset of S2 (comparison duration) and another one time looked to the onset of the participants’ response. The duration of all events was set to zero. All events were then convolved to the canonical hemodynamic response function (HRF). The linear models included also motion correction parameters as effects of no interest. The data were high-pass filtered with a cutoff frequency set to 0.0083 Hz. The 18 runs were then concatenated to avoid any deformation of the time-series due to the filtering^42^.

Our first aim was to identify brain regions responsive to all four S1 durations. To do so for each subject, we estimated 4 t-contrasts, one for each S1 duration, averaging across the 18 runs. The resulted contrast images (t-statistics maps in DARTEL space) were then used in a second-level ANOVA where, once again, 4 different contrasts were calculated. The statistical threshold was set to p < 0.05 FWE cluster-level corrected for multiple comparisons across the entire brain volume (cluster size estimated at a voxel level threshold p-uncorrected = 0.001). Correction for non-sphericity^43^ was used to account for possible differences in error variance across conditions and any non-independent error terms for the repeated-measures.

The 4 group-level t-maps were finally used to perform a “winner-takes-all” classification procedure, in which each voxel according to its t value was classified as maximally responsive to one of the four different S1 durations (an arbitrary value from 1 to 4 was assign to these voxels). All voxels with a t-value lower than 3.13 were excluded from this classification procedure (i.e., their value was set to 0).

#### First constrain

We considered regions of interest (ROIs), brain areas that had all four duration selective clusters (i.e., which included all 4 voxel-types) with a cluster size of at least 20 voxels per S1 duration. According to the above criterion, we identified 5 distinct ROIs: Cerebellum (Vermis VI, left lobule VI, right lobule V), left & right primary visual cortex (V1), left Intraparietal Sulcus (IPS), left Supplementary Motor Area (SMA) and left Inferior Frontal Gyrus (IFG, see Figure 1a and Supplementary Table 1 for more details).

#### Second constrain

Moreover, we considered ROIs only the regions in which the duration selective clusters of voxels show a tuning profile. The averaged (over voxels) hemodynamic response of each duration selective cluster was highest for the preferred duration (PD), as expected, and slowly decayed with durations distant from it (PD ±1 and 2, see Figure 2c).

The normalization was performed as follow:

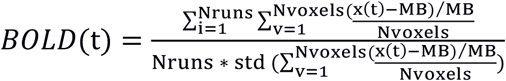

where t is the signal in a given voxel and MB is the baseline obtained averaging the signal of t across runs. Normalization was then performed subtracting the signal in a given voxel from a baseline value and dividing it by the baseline. The BOLD response was aligned to the 2^nd^ volume (i.e., a TR) after the offset of the S1 duration. Within a single subject, we first averaged the BOLD signal across the voxels of a cluster and then across the fMRI runs.

The second-level analysis on DARTEL normalized images, was performed only to identify the 5 ROIs. All the following connectivity analyses were performed in the individual subjects on images in the subjects’ native space.

### Mask Creation

After identifying the five ROIs at group-level (DARTEL space), we identified the same regions in each individual subject by using transformation matrices available in different stereotaxic atlases. For the creation of cerebellum mask, we employed SUIT (Spatially unbiased atlas template of the cerebellum and brainstem, http://www.diedrichsenlab.org/imaging/suit.htm) toolbox^44,45^. Specifically, we identify in each subject the following areas: Vermis VI, right lobule V and left lobule VI. For V1, we used the Destrieux cortical parcellation atlas available in FreeSurfer (image analysis suite, which is documented and freely available for download online at http://surfer.nmr.mgh.harvard.edu) ^46,47^ and we combined V1 and S-calcarine (left and right) labels. For IPS, SMA and IFG we used the Freesurfer’s Desikan-Killany cortical parcellation atlas^48^. Specifically, for IPS we used the intraparietal and parietal transverse sulcus labels. For SMA we used the medial part of BA6 label, while for IFG we combined the BA45 and BA 44 labels.

These masks were used later to identify the volumes of interest (VOIs); i.e. voxels within the ROIs that were significantly active for the contrasts of interest (i.e., 4 t-contrasts, one for each S1 duration).

### Dynamic Causal Modelling (DCM) analyses

DCM is a direct measure of Effective Connectivity^4,49^. The main objective of DCM is to estimate the coupling parameters of a hypothetical underlying neuronal model and evaluate how well the modeled BOLD signal approximate the observed BOLD response. These coupling parameters concern both the architecture of the network (A-matrix) and the influence of a given stimulus (i.e., experimental manipulation) on the strength of the connections (B-matrix) as well as on the activity of the network nodes (C-matrix). The term architecture refers to the summation of all possible connections between regions, which would imply a communication between them.

To put this explicitly, we modeled BOLD responses using the following DCM state equation:

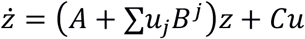

Where *z* is the BOLD signal, *u* represents the presented stimulus duration (i.e., S1 stimulus) j is the type of S1 (e.g. 0.4 s). A, B, and C are the different aspects of connectivity (i.e., network architecture, modulation of presented S1 on the existing connectivity and network nodes activity)

Here, we must stress that the number of possible combinations of A, B, C-matrices grows exponentially as the number of brain regions in the system grows. For small systems, investigating all possible connectivity architectures might be plausible, but this becomes unfeasible for large-scale systems. For example, in a network with 20 nodes we would have 2^20×20^ connections. Recent studies have tried to address this issue by either dividing the possible models into families^8^ or proceeding with Parametric Empirical Bayes (PEB) analysis^50^. DCM analysis was performed using DCM12 in SPM12 (v. 7219).

#### DCM and Parametric Empirical Bayes (PEB) of a 5-nodes network

Since we did not have a clear hypothesis about the network architecture of the 5 ROIs, we decided to use Parametric Empirical Bayes (PEB) which is considered a data-driven type of analysis.

First, for each subject, each fMRI run and each ROI we extracted the principle eigenvariate of the active voxels within the ROI i.e., voxels active at the offset of any S1 duration. In total, we ended up with 18 concatenated runs of 5 time-series (one for each ROIs), which were adjusted to the effects of interest (the five regressors of the first level analysis). Since we lacked of knowledge about the network connectivity in a temporal discrimination task, we decided to opt for the simplest model assumptions. So, we set the neuronal model of our DCMs to be deterministic, one-state, bilinear with mean-centered inputs. At this point is worth emphasizing that since the one-state model assumes excitatory activity only, we cannot make any inference about the type of neural activity (i.e., excitatory or/and inhibitory) contributing to the connectivity results.

To consider the fact that we acquired the images with a 7T MRI scanner, we revised the parameters of the BOLD signal equation^51,52,5^. Thus, in the file called “spm_gx_fmri.m” we set: EPI echo time at 0.026 s, ratio of intra- to extra-vascular signal at 0.026, intravascular relaxation rate at 340, frequency offset at 197.9 Hz and resting oxygen extraction fraction at 0.34. These values were changed according to the study by Heinzle et. al (2016)^52^.

Second, we specified a fully connected model between the five ROIs and we ran the DCM routine of 198 models (a model for each fMRI run and each subject i.e., 11 subjects x 18 runs) to calculate the parameters of the A-B-C-matrices best explaining the fMRI time-series.

Third, we used the PEB to find out the most probable network structure (i.e., strength of connections) across the 198 models^50^. Briefly, PEB seeks the optimal model by testing various A-B-C-structures and by switching off certain parameters (e.g. connections), it measures how this affects the model evidence. There are three main criteria which approximate the model evidence by identifying the optimum balance between accuracy (proximity to the observed BOLD signal) and complexity of a given model^53^. Among them, negative Free energy (F-criterion) is supposed to provide better Laplace approximation for the complexity term. This happens because the negative Free energy considers the interdependence between the estimated parameters while ensuring that they are as precise and as uncorrelated as possible^9,7,54.^ Models were treated as random effects that could differ between subjects. Random-effect analysis (RFX) is considered better when dealing with an unknown population distribution. It is also considered more robust to possible outliers^7^. Posterior probability for second-level effect was set at Pp>0.99.

After the identification of the optimal model across subjects with PEB, we proceeded with Bayesian Model Averaging (BMA), i.e. averaging the parameters across the “winning” models of all subjects. These parameters reflect the strength with which the activation changes in one network node affect the connected nodes. Positive values indicate facilitation of activation while negative values suppression of it^54^. Posterior probability for second-level effect was set at Pp>0.99.

#### DCM of a 20-nodes network

To explore the effective connectivity between the duration selective clusters we then ran a DCM on a 20-nodes network. The 20 nodes were the four duration selective clusters of voxels within the five ROIs (Cerebellum, V1, IPS, SMA, IFG). First, we extracted the fMRI time-course. For each subject and each fMRI run we took the principal eigenvariate for each duration-selective cluster of voxels within the ROI (the voxels active at the offset of the different S1 duration i.e., 0.2, 0.4, 0.6, 1.1 s). In total, we had 18 concatenated runs of 20 time-series (i.e., 20 VOIs), which were adjusted to the effects of interest (i.e., the five regressors of the first level analysis). We were unable to create the 20 VOIs for all participants, since not all of them show activation above threshold for each S1 duration within the 5 ROIs. The 20 nodes DCM was then performed only on 6 out of the 11 initial subjects. Although some statistical power was lost, the fact that we had 18 sessions per subject resulted in modeling 108 sets of time-series (i.e., 6 subjects by 18 sessions). This gave us enough statistical power to make inferences afterwards. After extracting the time-series, we set the neuronal model of our DCMs to be deterministic, one-state, bilinear with mean-centered inputs. As in the previous DCM we revised the parameters of the BOLD signal equation to be appropriate for a 7T MRI scanner.

Then, to simplify the complexity of the 20-nodes network, we incorporated the connectivity results (A-matrix) of the PEB analysis. More precisely, we allowed bi-directional connections between the 4 duration selective clusters of the Cerebellum to those of V1 and IFG, feedforward connections between Cerebellum and IPS, Cerebellum and SMA, V1 and SMA, IPS and SMA, and finally feedback connections from IFG to V1.

Since the model of connectivity between the five ROIs was borrowed from the PEB analysis, before testing specific hypotheses about connections and modulations by stimulus presentation in a 20-nodes network, we decided to do an extra check on the robustness of the PEB result. To do this, we ran 2 DCMs to determine the A-matrix best explaining the fMRI time-series. We compared two models one with an A-matrix “fully-connected” and one with a “PEB-like” structure. In both models B and C matrices were left fully-modulated. We then compared the two models by using Bayesian Model Selection (BMS). BMS is hierarchical and variational Bayesian approach which is more robust in dealing with outliers. This is because each model is treated as a random variable^7^. To measure the group evidence of the two models (“fully connected” versus “PEB-like”), we computed their protected exceedance probabilities (PEPs). PEPs are an improved extension of exceedance probabilities (Eps) in the sense that PEPs consider the possibility that the observed differences in model evidences (across subjects) might be due to chance^55^. EPs represent the belief that a given model is more likely than any other model, given the group data. We considered mandatory for a “winning” model to have PEPs> 90%.

Once we established that a “PEB-like” structure of the A-matrix was better than a “fully connected” one, we proceed by exploring which region among the five ROIs was modulated by S1 duration presentation (C-matrix). Fifteen possible models were tested. All models had an A-matrix “PEB-like”, a B-matrix in which all existing connections were modulated by stimulus presentation and 15 different combinations of C-matrices. The 15 combinations are the result of the fact that we arbitrarily allowed to have one or maximum two brain regions modulated by stimulus presentation. These 15 possible model-variations (i.e., nested models) were constructed using Bayesian Model Reduction (BMR). BMR is a rather new procedure that allows the computation of model evidence and posterior probabilities when the priors of a model with more parameters are known^50^. Bayesian Model Selection (BMS) showed that the most likely model was the one with a C-matrix in which IPS was the area modulated by duration presentation. For the winning model PEP was indeed greater than 90%. In BMS, models were treated as random effects.

At this point given a certain model structure (A-matrix “PEB-like”) and a C matrix in which IPS was the area modulated by duration presentation, we asked how the different duration-selective clusters of voxels communicate with each other (i.e., what is their connection and how these connections are modulate by duration presentation). To achieve this goal, we ran a DCM analysis, in which we tested hypotheses concerning the existence of connections (A-matrix “PEB-like”) and the modulation exerted by stimulus duration on both existing connections (B-matrix) and IPS activity (C-matrix). Specifically, we hypothesize these connections and modulations to be: 1) stimulus *duration independent*, 2) only partially stimulus duration dependent i.e., *neighboring dependent* or 3) totally stimulus *duration dependent*.

We tested 108 possible models which were the combination of these 3 possibilities (*i*.*e*., *duration independent, neighboring dependent, duration dependent*) applied to the A-B and C matrices (see equation below). Connections (A-matrix) and modulations (B-matrix) were checked within duration selective clusters on a given ROIs (w) or between duration selective clusters in different ROIs (b).

Ab^i^Aw^j^Bb^k^Bw^m^C^n^

Where ABC are the A-B-C matrices of the DCM analysis, i=1:3, j=1:3, k=i:3, m=j:3 and n=1:3, w is within ROI b is between ROIs.

To simplify our models, modulations of self-connections within each duration-selective cluster were not allowed (i.e., they were set to zero)^56^.

For the calculation of the A-B-C-parameters of these 108 models we used the BMR procedure. To identify the winning model, we first divided all models into three main families. The three families differed based on the modulatory effect of S1 stimulus duration on the activity of IPS duration selective clusters (i.e., stimulus *duration independent, neighboring dependent*, stimulus *duration dependent*). Each family had 36 models (i.e., 36 by 3 possible C modulations equal to 108 models) The PEPs of each family and model was then estimated using BMS, where models were treated as random effects^8,9^. To determine which of the 36 models within the winning family best explained our data, we used Bayesian Model Comparison (BMC)^9^. Finally, after the identification of the optimal model across subjects, we proceeded with Bayesian Parameter Averaging (BPA), i.e. we averaged all winning’s model A-B-C-parameters across subjects^10^.

## Acknowledgments

Financial support has been provided by the European Research Council -ERC (Grant Agreement No 682117, BiT-ERC-2015-CoG to D.B.), the Japan Science and Technology Agency (JST to R.K) and the Japan Society for the Promotion of Science (Grant Numbers JP17K20006, JP18H01101 and JP19H05313 to M.J.H). We would like to thank Dr. Mayur Narsude for supporting data collection.

## Author Contributions

FP and DB conceived the study; MJH collected the data; FP analyzed the data; FP, MJH, RK and DB wrote the paper.

