## Supplementary Material for "Topographic Connectivity in a Duration Selective Cortico-Cerebellar Network"

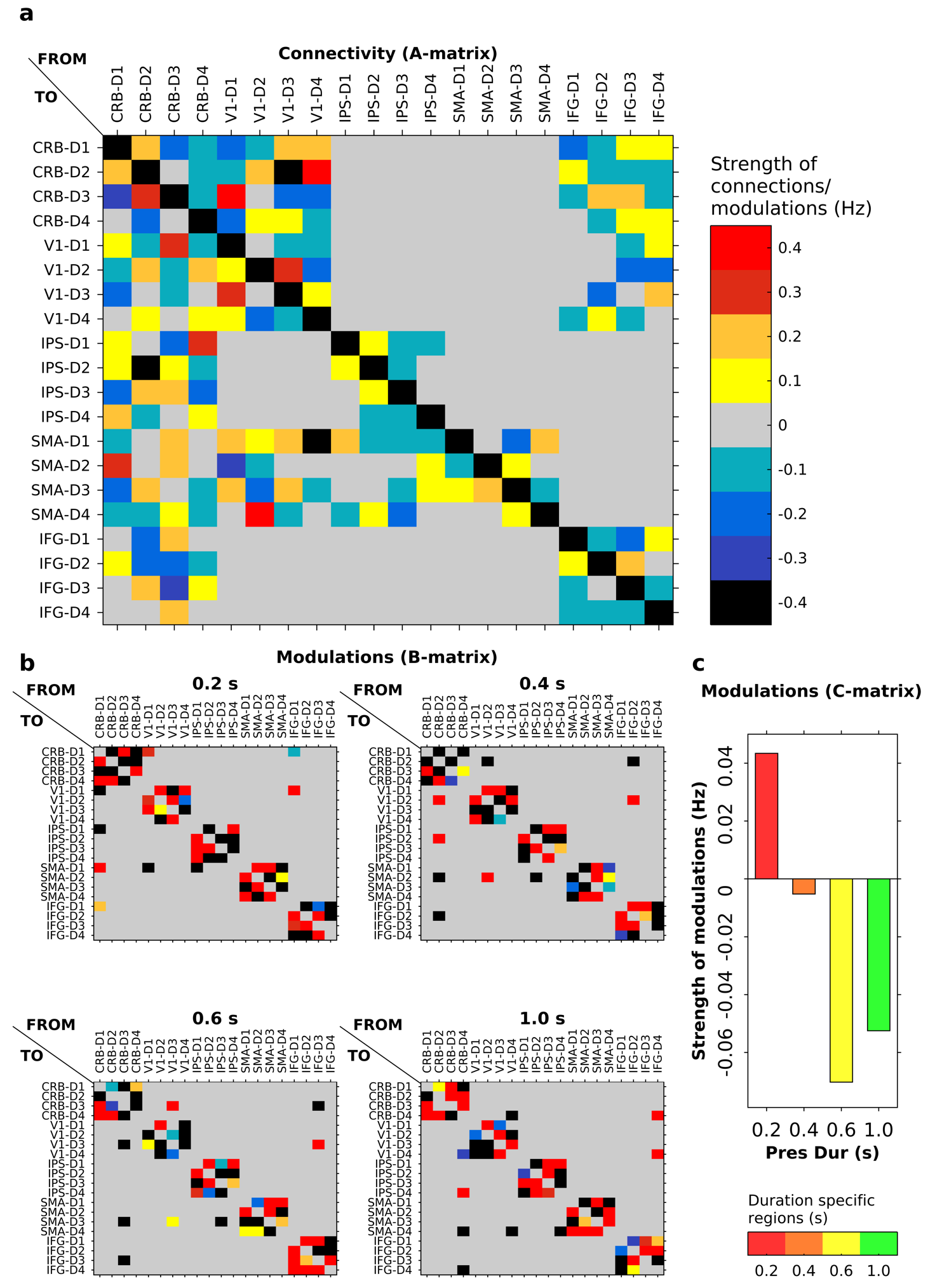

**Supplementary Figure 1 Bayesian model averaging (BMA) results of the winning model.** BMA averages the A-B-C-parameters of the winning model across subjects and sessions. (a) The matrix represents the strength of connections (A-matrix) between duration selective clusters of voxels. The color scale represents the parameter’s values. These values range from -0.4 to 0.4 Hz. (b) The four matrices represent the modulation on the connectivity strength (B-matrix) between the different duration selective clusters at the offset of the four S1 durations. x and y-axes are as in (a). (c) Bar-plot of the activity within each IPS duration selective cluster (C-matrix). Each bar shows the neuronal change (y-axis) for each of the four durations specific clusters while a specific duration was presented (x-axis). Duration specific clusters and stimulus durations are color coded as follow: red= 0.2, orange=0.4, yellow=0.6, and green= 1 s.

| **Anatomical Regions** | **DARTEL coordinates (mm)** | | | **Cluster size** | **p_FWE_** | **T** |
| --- | --- | --- | --- | --- | --- | --- |
| Left Inferior Parietal Lobule | -40 | -40 | 40 | 2524 | < 0.001 | 7.93 |
| Left Supplementary Motor Area | -2 | 9 | 46 | 699 | < 0.001 | 7.50 |
| Right Mid-Occipital Gyrus | 21 | -98 | 12 | 152 | 0.002 | 6.63 |
| Right Lateral Cerebellum | 32 | -51 | -20 | 433 | < 0.001 | 6.43 |
| Left Inferior Frontal Gyrus | -50 | 4 | 10 | 736 | < 0.001 | 6.42 |
| Right Cerebellar Vermis | 9 | -72 | -14 | 305 | < 0.001 | 5.89 |
| Right Lingual Gyrus | 18 | -45 | 2 | 264 | < 0.001 | 5.86 |
| Right Frontal Eye Fields | 27 | -6 | 52 | 122 | 0.009 | 5.59 |
| Left Calcarine Sulcus | -14 | -64 | 15 | 242 | < 0.001 | 5.42 |
| Right Middle Occipital Gyrus | 48 | -69 | 8 | 125 | 0.008 | 5.13 |
| Right Intraparietal Sulcus | 44 | -36 | 50 | 156 | 0.002 | 5.10 |
| Left Frontal Eye Fields | -24 | -6 | 46 | 219 | < 0.001 | 4.98 |

**Supplementary Table 1**

Stereotaxic Dartel-11 coordinates (mm) for brain areas activated at the offset of the four S1 durations. Voxels activated at p<0.05, FWE cluster-level corrected for multiple comparisons across the whole brain volume.

| **Networks** | **Analysis** | **A-matrix** | **B-matrix** | **C-matrix** | **N models** |
| --- | --- | --- | --- | --- | --- |
| **5-nodes** | PEB | all | all | all | - |
|  | DCM | all | all | all | 1 |
|  | DCM | PEB-like | all | all | 1 |
|  | DCM | PEB-like | all | 1 or 2 ROIs | 15 |
| **20-nodes** | DCM | PEB-like  I,N,D | I,N,D | I,N,D | 108 |
|  | DCM | PEB-like  I,N,D | I,N,D | D | 36 |

**Supplementary Table 2** Summary of the analyses performed.

Legend: PEB = Parametric Empirical Bayesian, DCM=Dynamic Causal Modelling, I=*duration independent*, N=*neighboring* *dependent*, D=*duration* *dependent*
